# Modeling the dynamic behaviors of the COPI vesicle formation regulators, the small GTPase Arf1 and its activating Sec7 guanine nucleotide exchange factor GBF1 on Golgi membranes

**DOI:** 10.1101/2020.09.06.285072

**Authors:** Garrett Sager, Ryoichi Kawai, John F. Presley, Elizabeth Sztul

## Abstract

The components and subprocesses underlying the formation of COPI-coated vesicles at the Golgi are well understood. The coating cascade is initiated after the small GTPase Arf1 is activated by the Sec7 domain-containing guanine nucleotide exchange factor GBF1. This causes a conformational shift within Arf1 that facilitates stable association of Arf1 with the membrane, a process required for subsequent recruitment of the COPI coat. Although we have an atomic level knowledge of Arf1 activation by Sec7 domain-containing GEFs, our understanding of the biophysical parameters that regulate Arf1 and GBF1 association with Golgi membranes and with each other is limited. We used Fluorescence Recovery After Photobleaching (FRAP) data and kinetic Monte Carlo simulation based on continuous-time random walk to assess behavior of Arf1 and GBF1 during COPI vesicle formation in live cells. Our analyses support a model in which Arf1 and GBF1 associate with Golgi membranes independently, with an excess of GBF1 relative to Arf1, and in which Arf1 activation is much faster than GBF1 cycling on the membrane. Interestingly, modeling the behavior of the GBF1/E794K mutant stabilized on the membrane is inconsistent with the formation of a stable complex between it and an endogenous Arf1, and suggests that its prolonged association with the membrane occurs independently of complex formation.

## INTRODUCTION

Membrane traffic is an essential process in eukaryotic cells, and is required to support such essential cellular activities as growth, compartment biogenesis and homeostasis, and sensing and responding to extracellular stimuli. Key events in membrane traffic are supported by a multitude of highly conserved genes, and deletion or mutation of such genes often lead to cellular death, underscoring the importance of the process (Howell et al., 2006; Olkkonen and Ikonen, 2006).

Membrane traffic within the secretory and endocytic pathways is mediated by vesicular carriers that form at the donor compartment, transit some distance through the cell, and then recognize and fuse with the acceptor membrane to deliver their cargo (Szul and Sztul, 2011; Wang et al., 2017). Vesicular traffic is selective and requires a mechanism to specifically package only some proteins into the forming vesicle, while excluding others (Derby and Gleeson, 2007; Tan and Gleeson, 2019). Cargo protein selection is mediated by a generally conserved mechanism that involves the assembly of a coat lattice on the cytosolic face of the donor membrane that “holds” cargo proteins within a spatially defined patch that subsequently invaginates to form a bud and ultimately pinches off as a vesicle. A family of structurally and functionally related coat complexes exist in cells to traffic proteins at different donor-acceptor membrane interfaces (Scales et al., 2000). At the ER-Golgi interface, vesicles coated with components of the COPII coat transport newly synthesized proteins from the ER to the Golgi, while vesicles coated with COPI components appear to retrieve escaped and cycling proteins from the Golgi back to the ER (Szul and Sztul, 2011).

The molecular events generating COPI vesicles from Golgi membranes have been reconstituted *in vitro* in a process requiring the activated GTP-bound form of the small Ras-related GTPase Arf1 and the heptameric coatomer complex (Ostermann et al., 1993; Spang et al., 1998). Like all GTPases, Arf1 cycles between the activated GTP-bound conformation and the inactive GDP-bound conformation. Arf1 is post-translationally modified through the attachment of the hydrophobic myristic acid at glycine in position 2 within its N-terminus, and this moiety is buried within the Arf1 while the protein is in the inactive GDP-bound form in the cytosol (Franco et al., 1995, 1996; Goldberg, 1998). To become activated, Arf1 must associate with a membrane and then be a substrate for an enzyme, a guanine nucleotide exchange factor (GEF) that facilitates the expulsion of the GDP and allows the binding of the activating GTP to the Arf.

Mammalian cells contain 15 Arf GEFs characterized by a highly conserved catalytic Sec7 domain, initially identified in the Sec7p protein of the yeast, *S. cerevisiae* (Bussey et al., 1983; Goldberg, 1998). The GEF required for Arf1 activation that leads to the formation of COPI vesicles has been identified as GBF1, a GEF belonging to the GBF1/BIG superfamily (Szul et al., 2005; Wright et al., 2014).

GBF1-catalyzed GDP/GTP exchange on Arf1 causes a conformational switch within Arf1 that exposes the hydrophilic myristic acid and allows its stable insertion into the membrane, securing the active Arf1-GTP on the lipid bilayer. Because of the allosteric coordination between the GDP/GTP exchange and myristic acid insertion, Arf1 activation can occur exclusively on the membrane. Furthermore, the insertion of the myristic acid is simultaneous with the alignment of the N-terminal α-helix of Arf1 onto the membrane, and the two processes confine Arf1 within a sterically restricted conformation on the membrane that exposes the effector-interacting interface (Nawrotek et al., 2016). The active Arf1-GTP then recruits cytosolic coatomer, but the exact stoichimetry of Arf1-coatomer binding is unclear, as multiple subunits of coatomer (β, γ, Δ and ε) contain Arf-binding sites (Szul and Sztul, 2011). Experimental evidence suggests that two Arf1 molecules, separated by 110 angstroms bind per coatomer (Yu et al., 2012). Repeated cycles of GBF1-mediated Arf1 activation and coatomer binding, combined with the recruitment of accessory proteins ultimately result in COPI vesicle formation.

Extensive enzymatic, biochemical and molecular analyses combined with crystallography, cryo-EM, and other structural approaches, provided an in depth understanding of how the Sec7 domains of GEFs mediate the Arf1-activating GDP/GTP exchange (Beraud-Dufour et al., 1998; Cherfils et al., 1998; Renault et al., 2002). However, we still are largely ignorant of how Arf1 and GBF1 diffuse within the cytosol to approach Golgi membranes, how they associate with Golgi membranes through transient and stable interactions, and how they interact with each other and other components once on the membrane. For example, GBF1 may have multiple roles during COPI coating, and in addition to binding and activating Arf1 to initiate the coating cascade, also binds the coatomer (Deng et al., 2009), perhaps “helping” Arf1 in recruiting the coat.

To determine the parameters of Arf1 and GBF1 behavior on Golgi membranes in live cells, we used FRAP and computational simulations under conditions where distinct subprocesses of Arf1 activation and/or coating were perturbed by molecular means. Significantly, while previous studies reported the FRAP of Arf1 and GBF1 when each protein was expressed individually in cells, we performed dual FRAP in cells co-expressing both proteins and imaged simultaneously. A comparison of single versus dual FRAPs shows a significant change in Arf1, but not GBF1 qualitative behavior. Our results suggest that Arf1 and GBF1 associate with Golgi membranes independently of each other, and that Arf1 dynamics are limited by reaction rate, while GBF1 is solely diffusion limited.

Our results also have more general implications for the way FRAP data are analyzed, especially when protein dynamics are measured during transient overexpression. Historically, the t_1/2_ has been used as the most popular metric to summarize FRAP data and to compare FRAP recoveries of different proteins. However, we now show that plotting FRAP data on a semi-log plot can highlight the qualitative features (diffusion or reaction(s)) that are most important for regulating a specific protein’s intracellular dynamics. Importantly, we also show that these qualitative features can be obscured when proteins are overexpressed in cells, suggesting that some phenotypes observed by studying the behavior of a single expressed protein may be an artefactual result of altering qualitative protein dynamics. Additionally, we document that studying the kinetic behavior of a wild-type protein when co-expressed with a mutant reaction partner can lead to a diminished phenotype caused by the reaction partner’s endogenous proteins.

## Results

### Coating sub-processes

We aim to describe parameters of Arf1 and GBF1 behavior on Golgi membranes during COPI vesicle formation. The overall coating process and the association/dissociation events that we are monitoring by FRAP and simulating are shown in Figure 1. In our model, Arf1-GDP and GBF1 arrive separately at Golgi membranes. Arf1-GDP has been proposed to bind to the SNARE membrin in a GTP-independent fashion prior to interacting with GBF1 (Honda et al., 2005). Arf1 and GBF1 subsequently form a complex (step 1), followed by the exchange of GTP for GDP and the insertion of Arf1 into the membrane (step 2). Arf1-GTP then separates from GBF1 while maintaining membrane association (step 3). Finally, Arf1-GTP forms a complex with coat and with GAP (step 4), which continuous formation of such complexes will ultimately lead to vesicle formation and its budding from Golgi membranes. To simplify the modeling scope, only Arf1 and GBF1 are explicitly considered, and all other reactions needed to form a vesicle are considered as one transition rate. To refine our models, we used molecular approaches to selectively perturb specific subprocesses through the expression of dominant inactive mutants of Arf1 and GBF1 (labeled underneath the membrane in Figure 1) and assess the effect of such changes on the FRAP of each protein.

**Figure 1:**
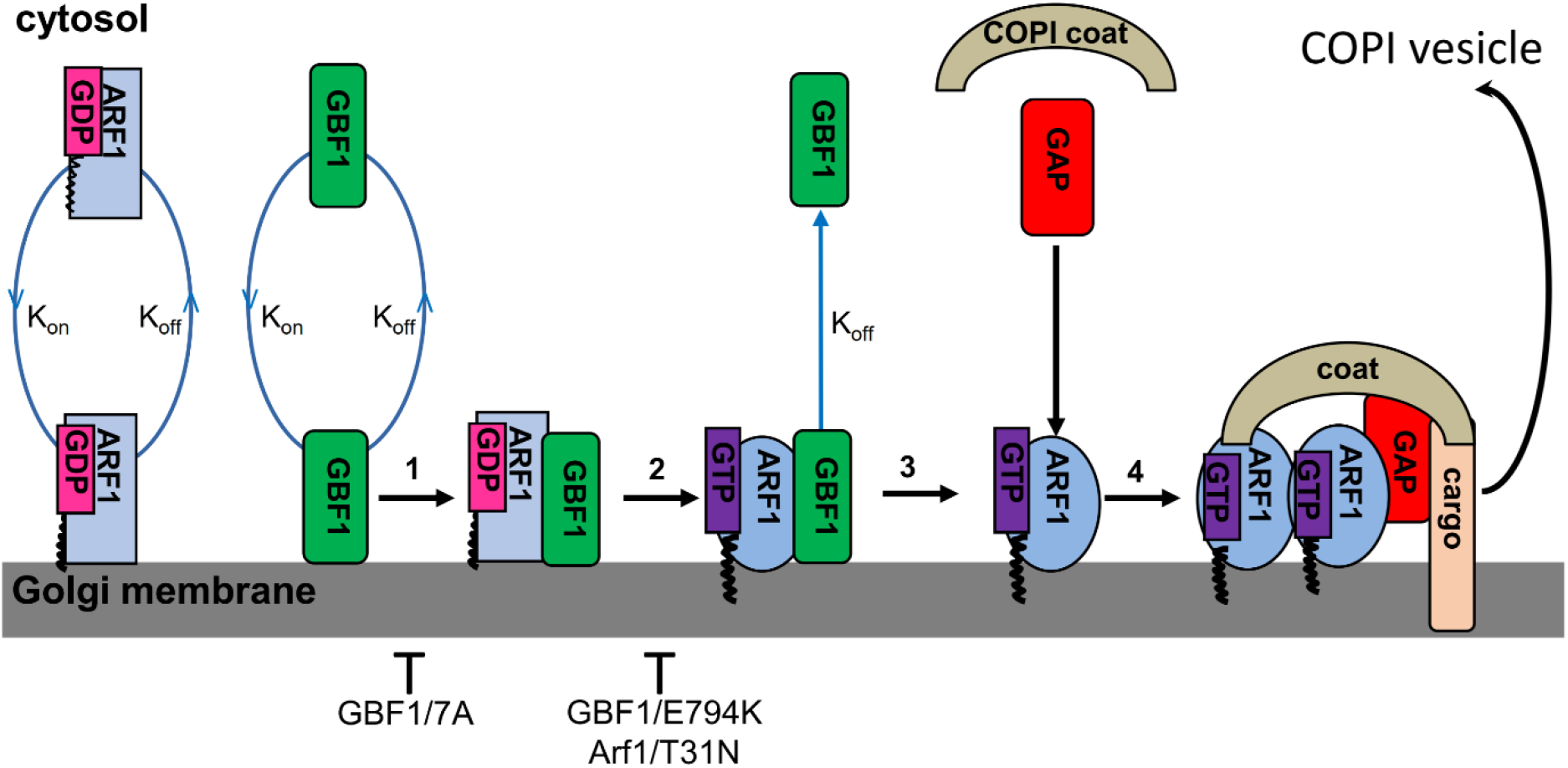
Subprocesses of COPI vesicle formation. A schematic of the 4 steps leading to Arf1 activation by GBF1 and the association/dissociation constants relevant for developing simulations for the dynamics of Arf1 and GBF1 during coating.

The contribution of the initial interaction of Arf1 and GBF1 to their membrane dynamics was probed by assessing the behavior of the GBF1/7A mutant and the behavior of Arf1 in cells expressing this mutant (Figure 1, step 1). The GBF1/7A mutant contains a mutation in a loop after helix J within its Sec7 domain, and although this GBF1 mutant associates with Golgi membranes, it is incapable of binding Arf1 (Lowery et al., 2011). When expressed in cells, GBF1/7A causes Golgi disruption, presumably by competing with the endogenous GBF1 for Golgi binding sites, thus preventing the activation of the endogenous Arf1. The importance of the next step, GDP expulsion from Arf1 by GBF1 to their dynamics was probed by examining the Fluorescence Recovery After Photobleach (FRAP) of the GBF1/E794K mutant and the behavior of Arf1 in cells expressing this mutant (Figure 1, step 2). GBF1/E794K was shown to associate with Golgi membranes and to bind Arf1, but is incapable of sterically displacing GDP from Arf1 (Garcia-Mata et al., 2003). The GBF1/E794K mutant also causes Golgi disruption, either by competing with the endogenous GBF1 for Golgi binding sites, or by competing for the Arf1 substrate, in both cases preventing Arf1 activation. We also perturbed the coating process by expressing the Arf1/T31N mutant previously shown to have low affinity for both GDP and GTP (Peters et al., 1995). Although we tentatively place its inhibitory action at step 2, it remains possible that Arf1/T31N does not stably associate with GBF1 (as explored below).

In all cases, we measured the FRAP of each protein when co-expressed with a cognate partner listed in Supplemental Table 1.

### Theoretical correlation between FRAP and reaction-diffusion kinetics

From a theoretical perspective, the exchange of a bleached protein or a protein complex on Golgi membrane measured by FRAP is generally either diffusion limited (restricted by how fast the protein diffuses in the cytosol and on the membrane), or reaction limited (regulated by how fast the protein associate/dissociate with the membrane and react with the relevant substrate or enzyme) (Sprague et al., 2004). Diffusion limited FRAP was the earliest analyzed case, and the solution for the diffusion constant, D, is well characterized

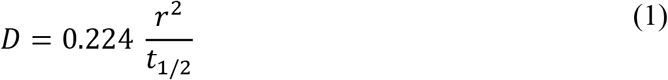

where the diffusion constant can be calculated by only knowing the time it takes for the photobleached region of interest (ROI) to reach half of its final intensity, t_1/2_, and the radius, r, of the bleached ROI (Axelrod et al., 1976; Edidin et al., 1976; Jacobson et al., 1976; Peters et al., 1974; Poo and Cone, 1974; Schlessinger et al., 1976; Zagyansky and Edidin, 1976). The t_1/2_ is a frequently used method to summarize FRAP data and allows direct comparisons of FRAP recoveries for multiple components within a single process (Nehls et al., 2000). These comparisons are generally interpreted as reflecting how fast or slow a protein recovers, and suggest which proteins may be “held back” by some additional process.

Most relevant for reaction limited FRAP data analysis is the effect of protein overexpression on reaction rates *via* the law of mass action. Conceptually, it is well agreed that overexpression can affect cellular dynamics through multiple mechanisms such as resource overload, stoichiometric imbalances, promiscuous interactions, and pathway modulation (Moriya, 2015). Resource overload occurs when the overexpressed protein requires resources which are needed by other pathways. For instance, if the protein uses chaperones, the overexpressed protein may be taking away chaperones needed to fold proteins required for a basic cellular function. Stoichiometric imbalance happens when a single subunit of a hetero-oligomer is overexpressed, which can cause an abnormal, toxic protein complexes (Abruzzi et al., 2002). Promiscuous interactions occur if an overexpressed protein contains a highly unstable, flexible region, since such domains can interact with low affinities with numerous proteins, leading to unpredictable consequences. Pathway modulation ensues when regulatory proteins are overexpressed, which may increase essential reaction rates and change the qualitative behavior of a pathway. While each of these four perturbation mechanisms seems conceptually probable, there is a lack of quantitative data assessing how much overexpression can change the qualitative interpretation of kinetic experiments. Here, we address this gap by using FRAP as a model kinetic experiment and the behavior of Arf1 and GBF1 during COPI vesicle formation at the Golgi as a model biological system.

Mathematically, determining whether a process is reaction or diffusion limited from FRAP data can be fairly simple. For a normalized FRAP curve of I(t) (intensity at time t), it can be achieved by plotting the inverse intensity, K(t) = 100 – I(t) (100 is used, because the FRAP data are normalized to 100). When the recovery is limited by one, first-order reaction, K(t) can be fit with a single exponential

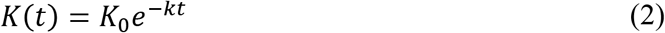

where *K*_0_ is 100 minus the mean intensity in the ROI in the frame immediately after the bleach and k is the reaction rate. Taking the natural logarithm of Eq. 1 yields

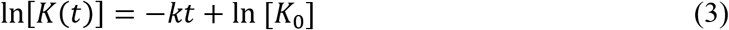

Comparing Eq. 3 to the linear equation, y=mx+b, shows the slope of Eq. 3 is −*k*. Plotting a hypothetical FRAP recovery limited by one, first-order reaction, as mathematically represented by Eq. 2, on a semi-log plot effectively takes Eq. 2 and makes function appear as Eq. 3. Therefore, the slope of the line on the semi-log plot is proportional to the negative reaction rate (Figure 2A). The more “downward” the line, the faster the reaction. FRAP data represented by a simple exponential decay indicates that the recovery is limited by a single rate process, such as on-off exchange between cytosol and membrane or a single reaction with another macromolecule. Diffusion is so fast that it is invisible in the hypothetical recovery curve.

**Figure 2:**
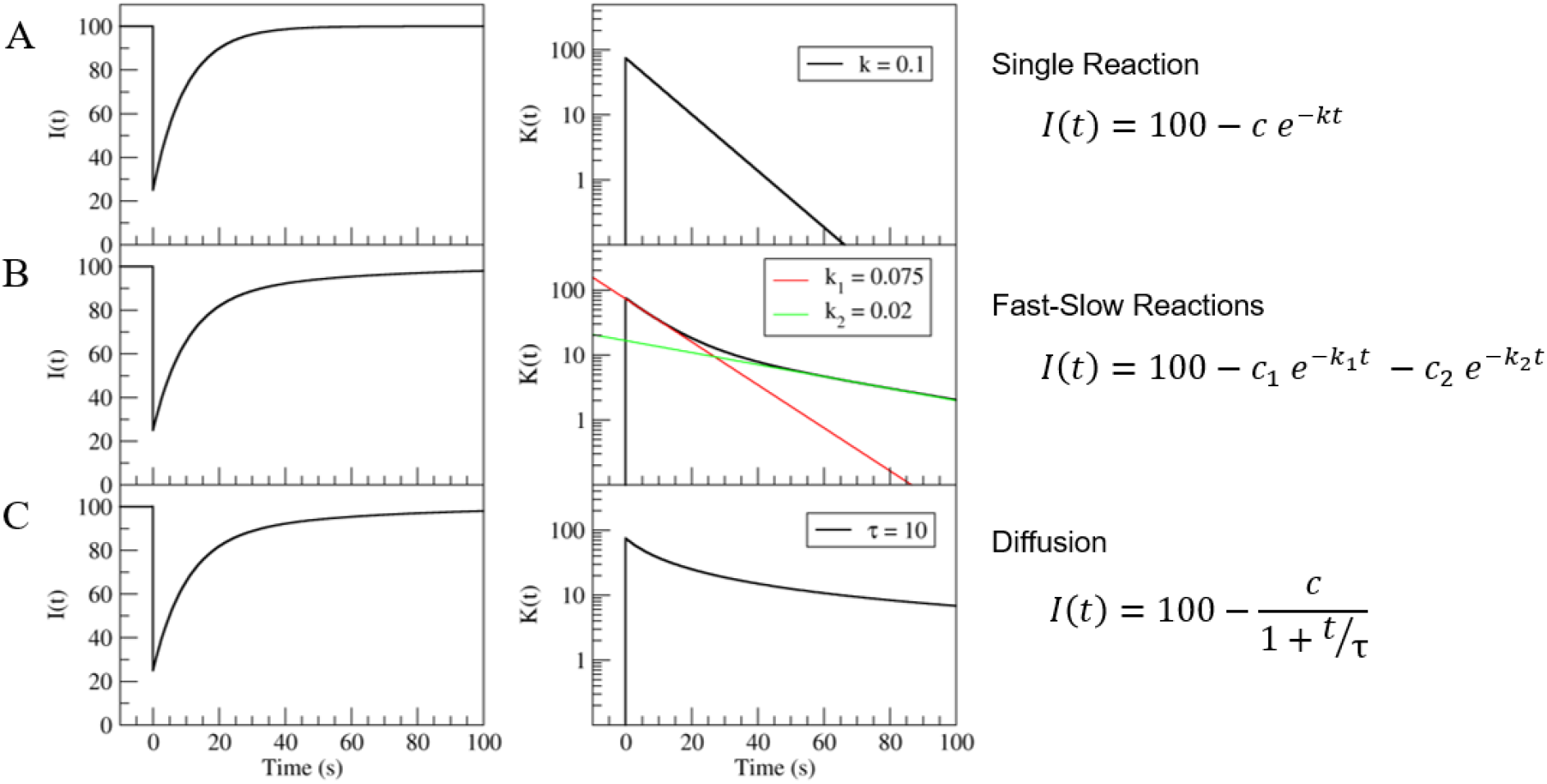
Theoretical plots of reaction-diffusion kinetic during FRAP. I(t) shows the normalized FRAP data on a conventional plot, while K(t) is 100-I(t) and is plotted on a semi log plot. Case A demonstrates one reaction rate, case B demonstrates two reaction rates, and case C represents diffusion.

When FRAP is limited by multiple first-order reactions, the data can be fit by a linear combination of exponentials (Figure 2B). In this example, a fast reaction dominates the dynamics for the initial part of the recovery (about the first 25 seconds), until the slower reaction limits the dynamics for the latter portion. In this scenario, when *K*(*t*) is graphed on a semi-log plot, the curve can be fit by two, straight lines. Unfortunately, it is difficult to visually differentiate between FRAP data limited by multiple reactions or diffusion (described below). In such cases, other information can help to make an educated prediction about whether the dynamic behavior of a protein is limited by multiple reactions or is diffusion limited. For example, knowing that a molecule has a small diffusion constant would suggest that its dynamics are likely limited by diffusion.

Another issue that can arise during FRAP analysis, especially when multiple reactions limit the protein’s kinetics, is the temporal resolution of the data. There is a tradeoff between sampling with high temporal resolution for a short time, or sampling with lower temporal resolution for longer time. For the semi-log method, we decided to use the former strategy. From past reports monitoring the recovery of COPI coat components over two or three minutes, it was observed that not much additional recovery occurs after 60 seconds, so there is little information to be gathered in the latter portion of the recovery curve (Bhatt et al., 2016; Niu et al., 2005; Szul et al., 2005). The first 30-40 seconds seem to be the richest in information. Thus, it is more advantageous to more densely sample the system for the first 40-60 seconds than done in previous reports. For instance, it is common for FRAP reports to measure the sample every 4-5 seconds. We chose to employ a higher sampling rate to allow detecting processes with shorter timescales.

The other common limiting process during FRAP is diffusion. In this scenario, K(t) can be fit by

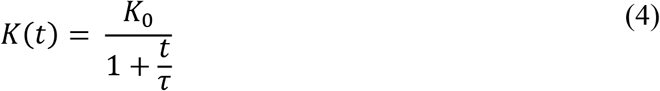

Where *τ* is a time constant that depends on the diffusion coefficient for the specific molecule. Taking the logarithm of Eq. 4 will generate a nonlinear curve (Figure 2C). However, it is important to note a non-linear function in the semi-log plot does not guarantee the molecular dynamics are limited by diffusion; it merely indicates that the molecular dynamics are not limited by a reaction. Using the same quantitative analysis, we used FRAP to understand the limiting processes of Arf1 and GBF1 diffusion and association with the Golgi membrane.

### Kinetic behavior of GBF1 is diffusion-limited

The FRAP behavior of wild-type GBF1 when expressed alone in cells is shown in Figure 3A, and is consistent with previously published values. As shown in Table 1, the t_1/2_ of 7.5 sec in this study is similar to the t_1/2_ of 9.4 sec reported previously in Bhatt et al., 2016, but lower than the previously reported values of t_1/2_ of 17 sec (Szul et al., 2005) and 30 sec (Niu et al., 2005). It is likely that the faster FRAP of the more recent studies is due to faster acquisition rates of now available imaging systems. Plotting the FRAP data on a semi-log produced a curve, suggesting that GBF1 recovery under these conditions might be diffusion limited.

**Figure 3:**
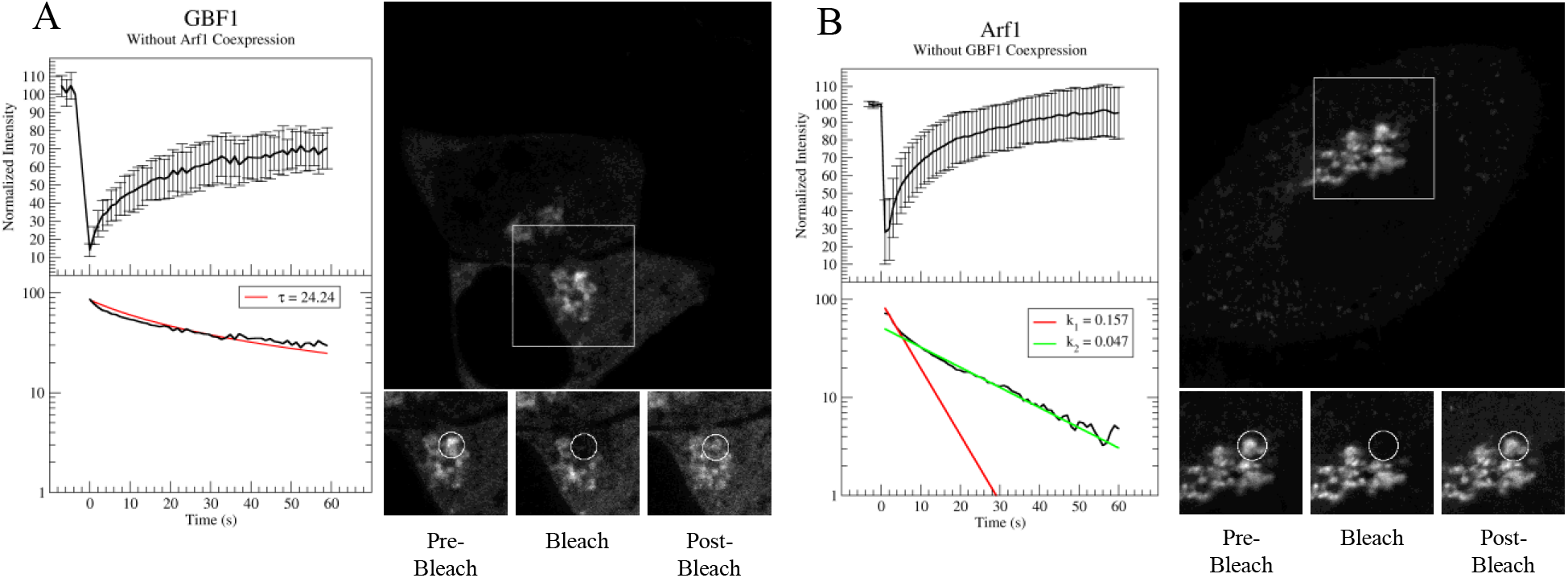
FRAP dynamics of GBF1 and Arf1 when expressed alone. HeLa cells expressing GFP-tagged wild-type GBF1 (A) or mCherry-tagged Arf1 (B) alone were processed using FRAP. The top plot in each panel shows the mean (n= 10) FRAP recovery with error bars representing the standard deviation (s.d.). The bottom plot in each panel shows the inverted mean FRAP data plotted on a semi-log plot. Representative pre-bleach, bleach, and post-bleach (after 54.6 seconds) images of the FRAP data are also displayed.

**Table 1.**
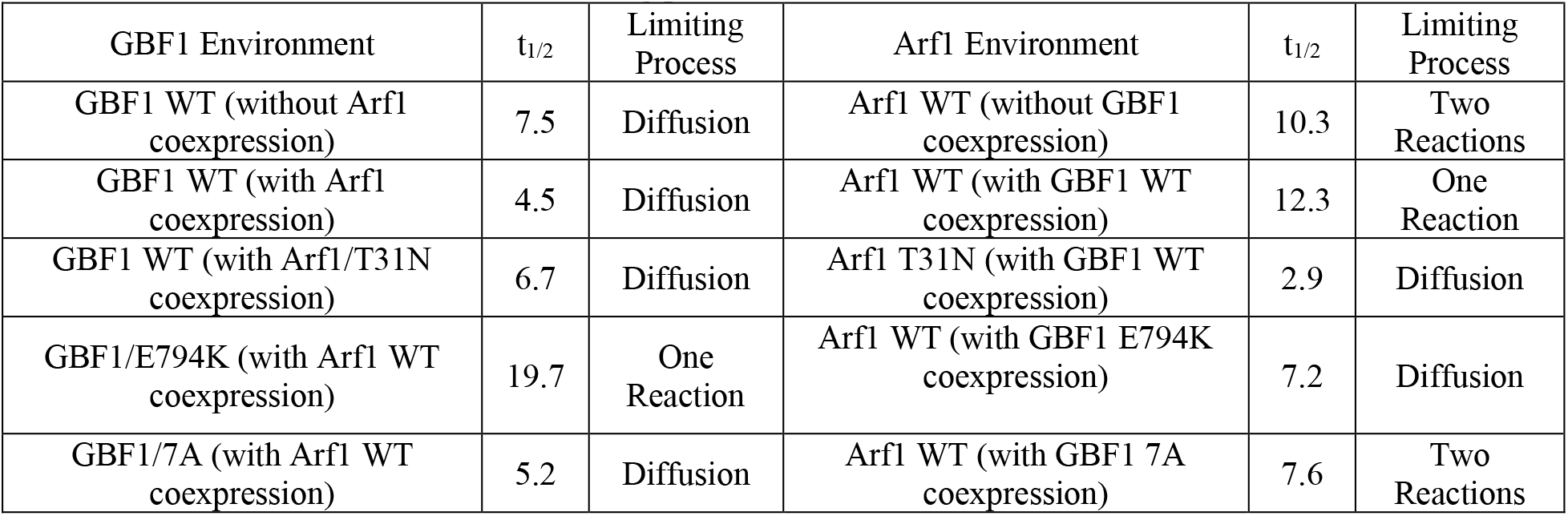
GBF1 and Arf1 FRAP t_1/2_ and limiting process under different conditions. The table lists the mean t_1/2_ values for GFP-tagged GBF1 constructs expressed alone or co-expressed with Arf1-mCherry constructs and for Arf1-mCherry alone or co-expressed with GFP-tagged GBF1 constructs. The standard deviation (STD) and statistical significance for these measurements are presented in supplemental Table 2 and 3.

| GBF1 Environment | $t_{1/2}$ | Limiting Process | Arf1 Environment | $t_{1/2}$ | Limiting Process |
| --- | --- | --- | --- | --- | --- |
| GBF1 WT (without Arf1 coexpression) | 7.5 | Diffusion | Arf1 WT (without GBF1 coexpression) | 10.3 | Two Reactions |
| GBF1 WT (with Arf1 coexpression) | 4.5 | Diffusion | Arf1 WT (with GBF1 WT coexpression) | 12.3 | One Reaction |
| GBF1 WT (with Arf1/T31N coexpression) | 6.7 | Diffusion | Arf1 T31N (with GBF1 WT coexpression) | 2.9 | Diffusion |
| GBF1/E794K (with Arf1 WT coexpression) | 19.7 | One Reaction | Arf1 WT (with GBF1 E794K coexpression) | 7.2 | Diffusion |
| GBF1/7A (with Arf1 WT coexpression) | 5.2 | Diffusion | Arf1 WT (with GBF1 7A coexpression) | 7.6 | Two Reactions |

The qualitative dynamics of GBF1 do not change when GBF1 is co-expressed with its Arf1 substrate, and plotting these FRAP data on a semi-log also produced a curve (Figure 4A and Table 1), suggesting that under these conditions, GBF1 recovery is also diffusion limited. These results imply that GBF1 dynamics on the membrane are predominantly defined by its diffusion rate, and indicates that the diffusion parameter is slower than any associations/reactions that GBF1 might have with the Arf1 substrate on the membrane. To test this hypothesis, we postulated and tested two key predictions.

**Figure 4:**
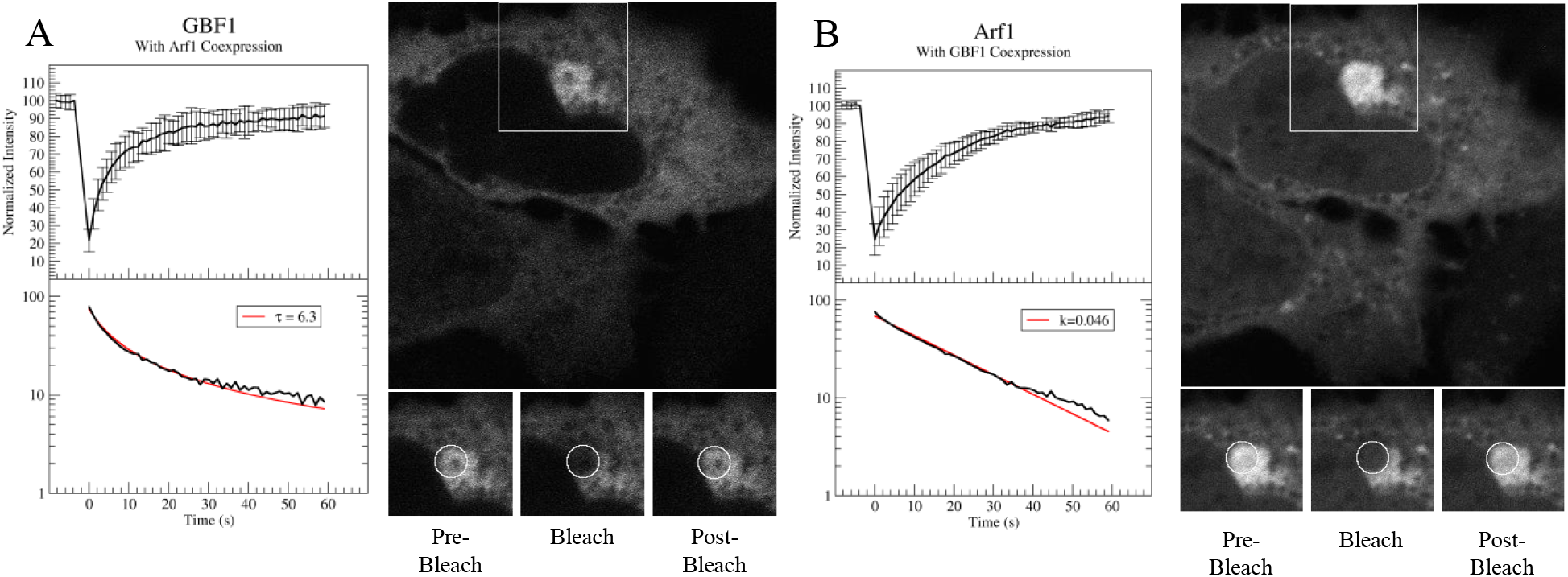
FRAP dynamics of GBF1 and Arf1 when co-expressed. HeLa cells co-expressing GFP-GBF1 and Arf1-mCherry were simultaneously imaged by FRAP. The top plot in each panel shows the mean (n= 11) recovery with error bars representing the standard deviation (s.d.). The bottom plot shows the inverted mean FRAP data plotted on a semi-log plot. Representative pre-bleach, bleach, and post-bleach (after 59.1 seconds) images of the FRAP data are also displayed.

First, since the dynamics of wild-type GBF1 appear not to be defined by its association/reaction with Arf1, the dynamics of a GBF1 mutant unable to bind Arf1 should parallel those of GBF1. We have previously generated and characterized the GBF1/7A mutant unable to bind the Arf1 substrate (Lowery et al., 2011). This mutant is unable to interact with the co-expressed Arf1 and its dynamics should be analogous to the dynamics of wild-type GBF1 expressed in cells without Arf1. Indeed, as shown in Figure 5A and Table 1, the t_1/2_ of GBF1/7A when co-expressed with Arf1 is similar to that of wild-type GBF1 (t_1/2_ of 5.2 sec for GBF1/7A when co-expressed with Arf1 versus t_1/2_ of 7.5 sec for GBF1 expressed alone). Plotting the FRAP of GBF1/7A on a semi-log results in a curve, suggesting that GBF1/7A behavior also is diffusion limited.

**Figure 5:**
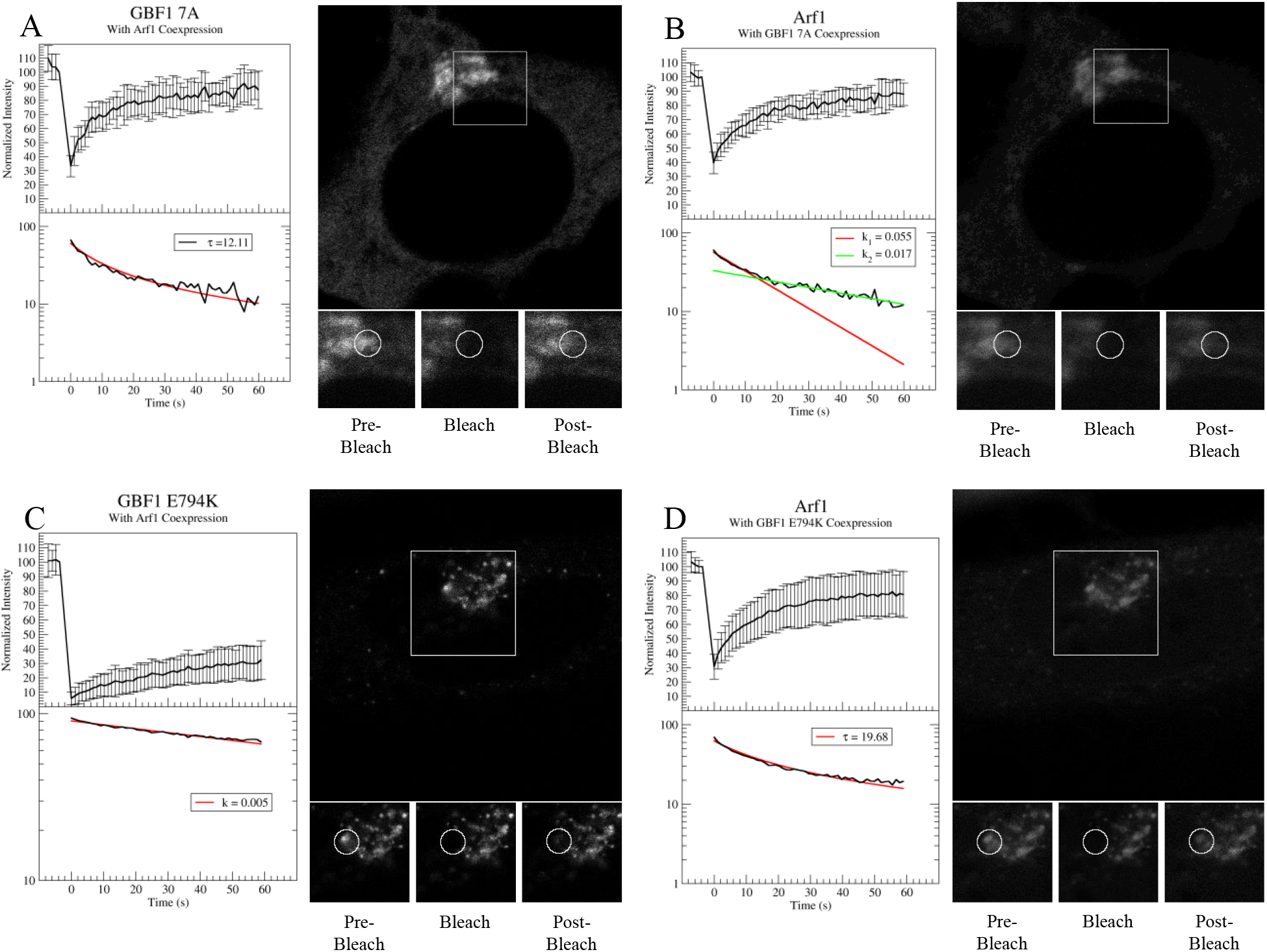
FRAP dynamics of GBF1 mutants and Arf1 when co-expressed. HeLa cells co-expressing GFP-GBF1/7A and Arf1-mCherry (A and B) or co-expressing GFP-GBF1/E794K and Arf1-mCherry were simultaneously processed using FRAP. The top plot in each panel shows the mean (n= 12) recovery with error bars representing standard deviation (s.d.). The bottom plot in each panel shows the inverted mean FRAP data plotted on a semi-log plot. Representative pre-bleach, bleach, and post-bleach (after 58.0 seconds) images of the FRAP data are also displayed.

Second, if the observed diffusion-limited FRAP behavior of wild-type GBF1 co-expressed with Arf reflects a slow diffusion parameter and a fast association/reaction with the Arf substrate (fast enough to be “invisible” within the FRAP dynamics), then the FRAP dynamics of the GBF1/E794K mutant would be expected to be more strongly influenced by the association/reaction component of the dynamics. GBF1/E794K contains a charge reversal from the acidic “glutamic finger” to a positively charge lysine in its Sec7 domain, and has been shown to bind Arf1, but doesn’t catalyze GDP expulsion from the bound Arf1. GBF1/E794K has been shown to be stabilized on membranes relative to wild-type GBF1 (Niu et al., 2005; Szul et al., 2005).

We examined the FRAP of GBF1/E794K co-expressed with Arf1, and show that it is significantly slower relative to wild-type GBF1 co-expressed with Arf1 (t_1/2_ of 19.7 sec versus t_1/2_ of 4.5)(Figure 5C and Table 1). When plotted on a semi-log scale, the membrane exchange dynamics of GBF1/E794K co-expressed with Arf1 is limited by one, slow reaction. This suggests that the membrane dynamics of GBF1/E794K are no longer regulated by diffusion, but instead are defined by membrane events. Previous reports suggested that the stabilization of GBF1/E794K on membranes reflects it remaining in a complex with the endogenous Arf1, with the implication that the expulsion of the GDP is the rate-limiting step in GBF1 on-off membrane dynamics. However, these results are also consistent with the possibility that GBF1/E794K is stabilized on the membrane in a process that is Arf1 independent (see below).

Together, our data are most consistent with a model in which the overall FRAP dynamics of wild-type GBF1 in cells reflect a fast association/reaction of GBF1 with Arf1 on the membrane. While the precise rate of this reaction cannot be resolved, it must be sufficiently fast that the overall reaction rate is limited by the slow diffusion of GBF1 within the cytosol.

### Qualitative dynamics of Arf1 are influenced by GBF1 co-expression

FRAP data for Arf1 when expressed in cells alone are shown in Figure 3B, and indicate that Arf1 recovery is limited by a fast reaction for the first few seconds, and subsequently by a slow reaction for the rest of the experiment. This suggests that Arf1 diffuses rapidly in the cytosol and its dynamics are predominantly regulated by two distinct membrane events, one extremely rapid and another with slower kinetics. A plausible model is that when Arf1 is expressed alone, it saturates the available pool of endogenous GBF1, and a proportion of the exogenously expressed Arf1 undergoes rapid unproductive cycles of association and dissociation from the membrane (the “fast” reaction), while the rest interacts with GBF1 on the membrane, is activated and participates in coating events (the “slow” reaction).

In contrast, when Arf1 is co-expressed with GBF1 (Figure 4B), Arf1 is also reaction limited, but the graph on semi-log indicates only a single reaction rate. This reaction rate appears similar to the “slow” reaction component observed for Arf1 when expressed alone. A plausible model is that when Arf1 and GBF1 are co-expressed, all the exogenously expressed Arf1 can interact with GBF1 productively and this is reflected in the kinetics, which are now limited by the one, potentially first-order, reaction.

Assuming that the co-expression situation reflects a more physiological relationship between levels of GBF1 and Arf1 in cells, our data suggest a model in which endogenous Arf1 would exhibit analogous dynamics and associate with Golgi membrane in productive interactions and activation through endogenous GBF1. In this model, there would be an excess of GBF1 on the membrane waiting for Arf1, so that all Arf1 molecules would be able to interact with GBF1 (only one slow reaction would be observed).

This model was tested by co-expressing Arf1 with GBF1/7A, a mutant of GBF1 that doesn’t bind Arf1 (Lowery et al., 2011). A prediction of our model would be that when co-expressed with GBF1/7A, Arf1 dynamics should be analogous to those observed when Arf1 is expressed alone. Indeed, as shown in Figure 5B, two reaction limited rates were detected, similar to Arf1 dynamics when expressed alone, supporting a model that without productive binding to GBF1/7A, Arf1 exists in a pool that unproductively interacts with the membrane and a pool that binds the endogenous GBF1 on the membrane.

We also examined FRAP of Arf1 when co-expressed with the enzymatically compromised GBF1/E794K mutant (Figure 5D). Based on the current model that GBF1/E794K is stabilized on membranes because it remains in an unproductive complex with Arf1, our expectation was that Arf1 dynamics will be analogous to the dynamics of GBF1/E794K and be limited by a single slow reaction rate. Surprisingly, plotting the FRAP graph of Arf1 co-expressed with GBF1/E794K on a semi-log indicates a diffusion limited process and is inconsistent with Arf1 stably interacting with GBF1/E794K on the membrane

We believe that the diffusion-limited dynamics of Arf1 co-expressed with GBF1/E794K may reflect the varied interactions that can occur between the exogenously expressed Arf1 (the species we follow by FRAP) and either the exogenously expressed GBF1/E794K or the endogenous wild-type GBF1. Because Arf1 dynamics do not correlate with the behavior of GBF1/E794K on the membrane, these findings suggests that the reaction-limited GBF1/E794K events on the membrane may occur through a mechanism independent of Arf1 (see below).

### Qualitative dynamics of Arf1/T31N suggest GEF-independent membrane association events

To gain additional insight into the regulatory paradigms of GBF1 and Arf1 membrane dynamics, we assessed the behavior of the Arf1/T31N mutant that contains a single amino acid substitution at the end of the G-1 motif, GX_4_GK(S/T) (Nuoffer and Balch, 1994). Work with the T27N mutant of Arf6 (analogous to the Arf1/T31N mutant) has shown that Arf6 makes direct contact with the bound GXP, and when mutated causes the loss of affinity for both GDP and GTP (Macia et al., 2004). However, *in vitro*, the Arf6/T27N mutant can be recovered with GDP, but can’t be loaded with GTP, suggesting that in cells it may exist either as a GDP-bound form, or as an apo-GTPase. Arf6/T27N is capable of binding its GEF EFA6 in cells (albeit the efficiency of such binding was not determined, (Macia et al., 2004)), and the dominant negative effects of Arf6/T27N in cells have been ascribed to Arf6/T27N interacting with the endogenous EFA6 and preventing the activation of endogenous Arf6. However, it also remains possible that Arf6/T27N causes dominant negative effects not by saturating the endogenous GEF, but by preventing membrane association of the endogenous Arf6 and thereby preventing its activation.

Expression of Arf1/T31N in cells causes the dissociation of the COPI coat and the disruption of Golgi architecture, presumably due to lack of activation of endogenous Arf1 (Nuoffer and Balch, 1994; Peters et al., 1995). As for Arf6/T27N, it has been postulated that Arf1/T31N causes its dominant negative effects by saturating the endogenous GBF1 and competing with the endogenous Arf1 for binding and activation.

To test this model, we compared the FRAP dynamics of GBF1 when co-expressed with wild-type Arf1 to those when GBF1 is co-expressed with Arf1/T31N. As shown in Figure 6A and Table I, the FRAP of GBF1 is slightly slower in cells expressing Arf1/T31N (t_1/2_ of 6.7 sec as compared to t_1/2_ of 4.5 when co-expressed with wild-type Arf1). A previous report described a significantly larger stabilization of GBF1 (an increase from a t_1/2_ of 17 sec to a t_1/2_ of 45.6 sec), but in that study the co-expression of the GFP-tagged GBF1 and Arf1/T31N was assumed based on the morphology od the Golgi, but not actually visualized (Szul et al., 2005). Additionally, choosing cells sufficiently bright to image for FRAP is a subjective process, which may vary between experimentalists. This variance might reflect differences in protein density at the Golgi, which this report shows is a key parameter regulating FRAP data and may further lead to discrepancies in the recorded t_1/2_ values. As far as we are aware, ours represents the first dual FRAP that measures the dynamics of both, GBF1 and Arf1/T31N in the same cell. To provide insight into the impact of Arf1/T31N on GBF1, we plotted the GBF1 FRAP data on a semi-log scale, obtaining a curve (Figure 6A). These results indicate a diffusion-limited behavior for GBF1, unlikely to be strongly regulated through reactions with Arf1/T31N on the membrane.

**Figure 6:**
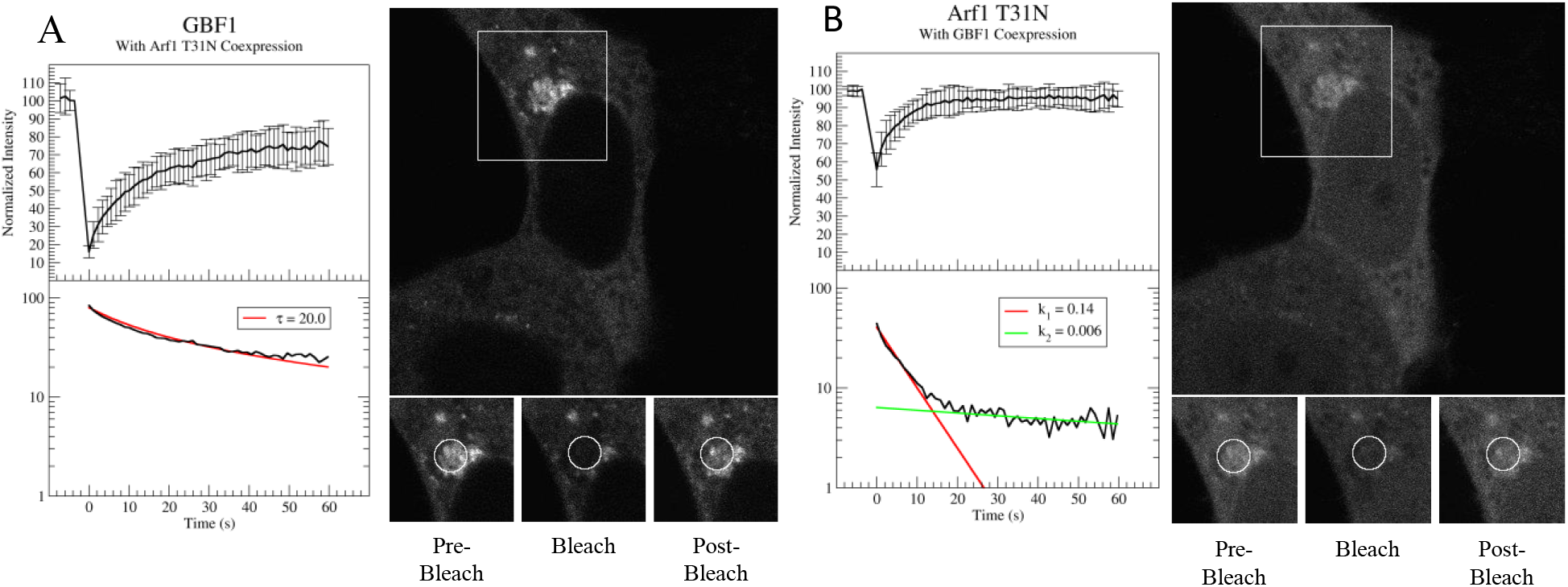
FRAP dynamics of GBF1 co-expressed with Arf1/T31N mutant. HeLa cells co-expressing GFP-GBF1 and Arf1/T31N-mCherry were simultaneously processed using FRAP. The top plot of each pnel shows the mean (n= 11) recovery with error bars representing standard deviation (s.d.). The bottom plot in each panel shows the inverted mean FRAP data plotted on a semi-log plot. Representative pre-bleach, bleach, and post-bleach (after 59.7 seconds) images of the FRAP data are also displayed.

This explanation was supported when we compared the FRAP dynamics of Arf1/T31N co-expressed with GBF1 with those of wild-type Arf1 co-expressed with GBF1. As shown in Figure 6B, a curve was obtained when FRAP data were plotted on a semi-log scale, suggesting that Arf1/T31N membrane dynamics are not limited by a reaction with GBF1. Thus, Arf1/T31 association/interaction with GBF1 on the membrane are so rapid as to be invisible within the cycling dynamics of Arf1/T31N.

Our findings are most consistent with a model in which Arf1/T31N causes its dominant negative effects in cells not by interacting with GBF1 and preventing GBF1 from activating endogenous GBF1, but rather from Arf1/T31N saturating membrane binding sites for endogenous Arf1. This model is supported by the observed intracellular localization of Arf1/T32N expressed in cells: Arf1/T31N can be detected in punctate structures where it co-localizes with GBF1, but the majority of Arf1/T31N appears to be diffusely distributed and not co-localize with GBF1 (Szul et al., 2005). Our model also is consistent with previous reports that Arf1 can associate with membranes prior to being activated by a GEF (Zhao et al., 2006).

### Simulation of GBF1 and Arf1 dynamics on Golgi membrane

To simulate the diffusion and reactions of GBF1 and Arf1, a simulation code based on kinetic Monte Carlo algorithm of discrete space, continuous time random walk was developed. The code is written in Fortran 95. We consider three molecular species GBF1, Arf1-GDP, and Arf1-GTP that are mobile. Their movements are simulated as jump processes with a fixed distance corresponding to the size of a typical protein. Erickson et al. provide several proteins of known mass (ranging from 40 kDa to 390 kDa) and dimensions as a reference, and these numbers were used to roughly estimate the dimensions of a standard protein as 4 nm x 4 nm x 10 nm (Erickson, 2009). The Arf1 diffusion constant has been calculated to be D=15 μm^2^/sec (Elsner et al., 2003), which is the number used in the simulation. The jumps are considered as Poison processes with a given mean transition time. The event driven algorithm based on a binary tree calendar is adopted for event scheduling to increase computational efficiency (Rapaport, 1995).

In total, there are 14 parameters for chemical dynamics, such as cytosol diffusion constant and membrane association/dissociation rates of Arf1-GTP (Table 2). The only parameter empirically obtained is the Arf1 diffusion constant in the cytosol (Elsner et al., 2003), and the other parameter values must be determined by fitting the simulation data to the experimental FRAP data. We analyzed 10 to 20 different parameter sets at a time, and initial guesses for the new sets of parameters were manually generated. Guesses were then refined by taking the new best fit and further guessing new set of parameters until there was a good qualitative agreement between the simulation and experiment. Every parameter, not experimentally determined, was fit independently for each experiment, unless indicated otherwise in the legend of Table 2. However, some parameters, such as the estimated diffusion constant for GBF1 were kept the same for each simulation. Analysis of parameters was repeated until a reasonable fit was obtained with FRAP of co-expressed wild-type GBF1 and Arf1 (Figure 7A).

**Table 2:** Simulation parameters for each analyzed species. The table lists the parameters used in the Monte Carlo simulation to generate the fit to the experimental data. The * denotes data gathered from published literature (Elsner et. al., 2003). The kon and koff values for Arf1-GTP are not necessarily indicative of the real membrane association and dissociation rates. These values were chosen to ensure that Arf1-GTP molecules remained on the membrane. The parameters that were allowed to vary from the wildtype simulation when trying to fit the mutant data are noted with the superscript †. The total number of molecules for GBF1, Arf1-GDP, and Arf1-GTP were allowed to vary for all mutant fits.

|  |  |  |  |  |
| --- | --- | --- | --- | --- |
| GBF1 species | WT | 7A | E794K | WT |
| Arf1 species | WT | WT | WT | T31N |
| GBF1 total molecules | 11239<sup>†</sup> | 9553<sup>†</sup> | 11239<sup>†</sup> | 4355<sup>†</sup> |
| GBF1 diffusion constant ($\mu\text{m}^2/\text{s}$) | 1.56 | 1.56 | 1.56 | 1.56 |
| GBF1 $k_{\text{on}}$ ($\mu\text{m}/\text{s}$) | 1.25 | 0.65<sup>†</sup> | 3.73<sup>†</sup> | 1.25 |
| GBF1 $k_{\text{off}}$ (1/s) | 1.58 | 1.58 | 0.43<sup>†</sup> | 1.58 |
| Arf1 GDP total molecules | 7031 | 7031 | 7031 | 12455 |
| Arf1 GDP* diffusion constant ($\mu\text{m}^2/\text{s}$) | 15* | 15* | 15* | 15* |
| Arf1 GDP $k_{\text{on}}$ ($\mu\text{m}/\text{s}$) | 12 | 12 | 12 | 12 |
| Arf1 GDP $k_{\text{off}}$ (1/s) | 0.18 | 0.18 | 0.18 | 0.39<sup>†</sup> |
| Arf1 GTP total molecules | 7296 | 7296 | 7296 | 12926 |
| Arf1 GTP *diffusion constant ($\mu\text{m}^2/\text{s}$) | 15* | 15* | 15* | 15* |
| Arf1 GTP $k_{\text{on}}$ ($\mu\text{m}/\text{s}$) | 187.5 | 187.5 | 187.5 | 187.5 |
| Arf1 GTP $k_{\text{off}}$ (1/s) | 0.05 | 0.05 | 0.05 | 0.05 |
| GBF1 / Arf1 association rate | 0.22 | 0.22 | 0.22 | 0.072 |
| GBF1 / Arf1 dissociation rate | 0.06 | 0.06 | 0.005<sup>†</sup> | 0.03<sup>†</sup> |
| Arf1 hydrolysis rate | 1.25 | 1.25 | 1.25 | 1.25 |

**Figure 7:**
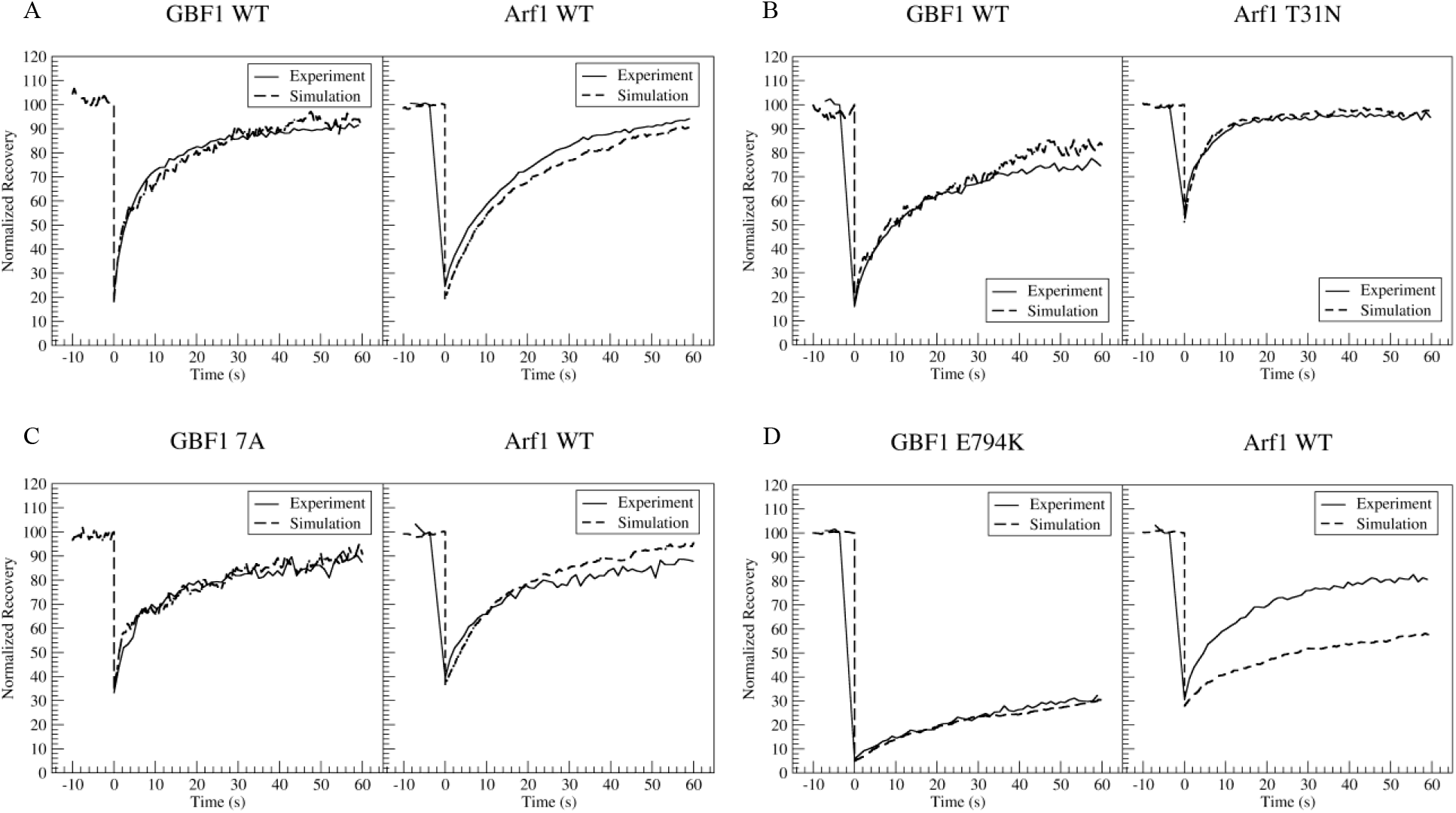
Simulation fits of FRAP data. HeLa cells co-expressing GFP-GBF1 and Arf1-mCherry (A) or GFP-GBF1 and Arf1/T31N-mCherry (B) or GFP-GBF1/7A and Arf1-mCherry (C) or GFP-GBF1/E794K GFP and Arf1-mCherry were imaged simultaneously using FRAP as described in Figures 4–6. In each panel, the solid line represents the experimental FRAP data, while the dotted line represents the best fit obtained by the simulation.

To test the simulations, we investigated how the parameter set changes to fit the FRAP data of GBF1 and the Arf1 when distinct subprocesses are perturbed by the expression of the Arf1/T31N, GBF1/7A or the GBF1/E794K mutant. An accurate fit is obtained for both proteins when GBF1 is co-expressed with Arf1/T31N (Figure 7B). Similarly, a good fit is evident for the GBF1/7A mutant co-expressed with wild-type Arf1, with only minimal disagreement with Arf1 FRAP at later recovery points (Figure 7C). In contrast, when GBF1/E794K was co-expressed with Arf1, the GBF1/E794K curve fits well to the experimental FRAP, but we could not obtain a good fit for Arf1 (Figure 7D). In our simulations, the theoretical Arf1 FRAP curve is much slower than the experimentally determined FRAP.

We can postulate a possible explanation for the faster than expected FRAP of Arf1 co-expressed with GBF1/E794K. In our experiments, the exogenously expressed GBF1 and Arf1 species are present in cells at approximately twice the level of the endogenous protein (data not shown). Thus, when GBF1/E794K and Arf1 are co-expressed, distinct interactions between the exogenously expressed and the endogenous proteins can occur. This variability is not problematic when assessing the FRAP of GBF1/E794K in cells co-expressing Arf1, since all GBF1/E794K molecules will interact exclusively with wild-type Arf1 (either exogenously expressed or endogenous). However, the situation changes when assessing Arf1 behavior in cells co-expressing GBF1/E794K, since the exogenous Arf1 (the species we follow by FRAP) can interact with two distinct species of GBF1, the exogenous GBF1/E794K mutant and the endogenous wild-type GBF1. Our simulation does not account for the presence and the contributions of the endogenous GBF1, and thus the theoretical FRAP of Arf1 shown in Figure 7D simulates Arf1 behavior in a cell depleted of endogenous GBF1 and expressing only the exogenous GBF1/E794K.

## DISCUSSION

Membrane traffic between the compartments of the early secretory pathway is mediated by anterograde vesicles coated with COPII components that transport cargo proteins from the site of their synthesis in the ER to the Golgi and by retrograde vesicles coated with the COPI coat that return membrane and cycling components from the Golgi to the ER (Szul and Sztul, 2011). The formation of COPI vesicles is initiated by the GBF1-mediated activation of Arf1 GTPase, which in its GTP-bound form recruits the heptameric coatomer to the membranes to initiate the assembly of the COPI coat. Importantly, both GBF1 and Arf1 are soluble proteins that associate peripherally with Golgi membranes and rapidly cycle between two pools – a larger cytosolic pool and a much smaller membrane-associated pool (Godi et al., 2004; Zhao et al., 2006). We are interested in understanding the parameters that regulate GBF1 and Arf1 membrane dynamics as means to gain insight into their association, interactions and functions in vesicle formation.

FRAP analyses have provided important insight into the dynamic exchange of peripheral membrane proteins that cycle between a cytosolic and membrane-associated pools, and we have used FRAP in conjunction with simulations to probe the relationships between GBF1 and Arf1 at the Golgi of live cells. FRAP is routinely used to compare the t_1/2_ of recovery of a particular protein when challenged with different conditions to generate models of its behavior and function. Here, we also used FRAP data to compare t_1/2_ times when distinct subprocesses of COPI coating were disrupted, but in addition, plotted the inverse of the FRAP measurements on a semi log plot to extracted more information about the behaviors of Arf1 and GBF1 under different cellular conditions. The semi-log plots inform on the type of behavior of a protein, as a reaction-limited FRAP is reduced to a straight line with a slope equivalent to the reaction rate, while pure and effective diffusion-limited FRAP is nonlinear on the semi-log plot.

This report presents a quantitative analysis of the behaviors of wild-type and mutant forms of GBF1 and Arf1 obtained from FRAP experiments on the Golgi in live cells under different conditions. Previous FRAP data have been used to propose certain behaviors and interactions between GBF1 and Arf1, but our analyses suggest that some of these models should be revisited.

### Implications for intracellular GBF1 dynamics

Previous FRAP studies of GBF1 expressed alone in cells reported a t_1/2_ of 30 sec (Niu et al., 2005) or 17 sec (Szul et al., 2005), and more recently 9.4 seconds (Bhatt et al., 2016). In this study, we obtained a t_1/2_ value of 7.5 seconds, and it is likely that the decrease in the t_1/2_ represents technological advances in photobleaching and the speed of image acquisition. FRAP of GBF1 expressed alone was nonlinear when plotted on a semi-log plot, suggesting that the process is diffusion-limited. This is consistent with the large size of the GBF1 dimer (~880kDa), as large proteins or proteins that exists in complexes are likely to experience steric hinderance during diffusion through the crowded cytoplasmic space packed with organelles and cytoskeletal elements (Saxton, 1993). The expression for the square distance, x, a molecule with diffusion coefficient, D, is expected to travel after some time, t, is

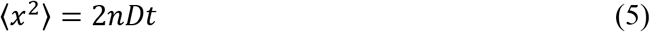

where n is the dimension of the brownian motion (n is 2 on the membrane, while n is 3 in the cytosol) (Michalet, 2011). The interpretation is that n describes the environment the molecule is navigating, and the time, t, is the typical time needed for a molecule with diffusion constant, D, to travel a distance of x^2^. Since D and x^2^ are directly proportional in Eq. 5, as D decreases, the distance the molecule is expected to travel in time must decrease. If this decrease is sufficiently large, the molecule’s dynamics will be limited by its small diffusion constant.

Such limited diffusion has been reported for the coatomer, a large complex of seven subunits whose measured diffusion coefficient was an order of magnitude slower than the predicted value calculated from the cross-sectional area of the coatomer (Elsner et al., 2003). The previously measured diffusion constant of the coatomer, using FRAP, is 1.7 μm / s^2^ (Elsner et al., 2003), which is strikingly similar to the diffusion constant determined for GBF1 by our simulation, 1.56 μm / s^2^. These values are consistent with the similar native sizes of the coatomer (650-700 kDa determined by filtration (Waters et al., 1991)) and of GBF1 (~880 kDa determined by Blue Native gels (Bhatt et al., 2016)).

To assess how interactions with Arf1 may affect GBF1 dynamics on the membrane, we examined FRAP of GBF1 when co-expressed with Arf1. A slightly quicker (but not significantly different: Supplemental Table 2) t_1/2_ of 4.5 sec was observed for GBF1 co-expressed with Arf1. We interpret the quicker cycling of GBF1 when co-expressed with Arf1 as suggestive that when expressed alone, the majority of GBF1 associates with membranes independently of Arf1 and remains associated for ~7.5 seconds while “looking” for Arf1, but when GBF1 is co-expressed with Arf1, GBF1 associates with membranes, rapidly finds membrane-associated Arf1, catalyzes GDP/GTP exchange on Arf1 and dissociates from the membrane. FRAP of GBF1 expressed singly or co-expressed with Arf1 is diffusion-limited, indicating that the processes of GBF1 binding and catalyzing Arf1 activation are significantly faster than GBF1 diffusion, and thus are not invisible on the semi-log plot.

Although the dynamics of GBF1 are not strongly influenced by Arf1 co-expression, it remains possible that other Arfs may have a stronger effect on GBF1 behavior. GBF1 is known to interact with Arf3, 4 and 5 when probed by bimolecular fluorescence complementation assay (Niu et al., 2005) and has been shown to activate those Arfs at the Golgi (Bhatt, 2019). In the future, examining GBF1 behavior when co-expressed with other class I and class II Arfs will allow more detailed analyses of its dynamics and behaviors.

Our co-expression FRAP studies suggest that both, GBF1 and Arf1 associate with Golgi membranes independently of each other, presumably by binding organelle-specific components. For GBF1, those could be proteins such as the Rab1 GTPase (Garcia et al., 2011) and the C10orf76 protein (Chan et al., 2019) or a specific phosphatidyl-inositol-phosphate (PI3P, PI4P and PI(4,5)P2) lipid components (Meissner et al., 2018). For Arf1, it could be the membrin SNARE (Honda et al., 2005) or hydrophobic interactions between the N-terminal a-helix and the membrane (Mouratou et al., 2005).

The dynamics of the GBF1/7A mutant unable to bind Arf1 co-expressed with Arf1 parallel those of wild-type GBF1 expressed alone (t_1/2_ of 5.2 sec versus t_1/2_ of 7.5 sec for wild-type GBF1), and in both cases, the GBF1 species is diffusion-limited. Previous work documented that the GBF1/7A mutant causes the dissociation of the COPI coat (a reflection of inhibition of Arf1 activation) and Golgi disruption (Lowery et al., 2011). Our FRAP analyses support a model in which GBF1/7A causes Golgi disassembly by competing for membrane binding sites with the endogenous GBF1 thereby preventing the activation of endogenous Arf1 by the endogenous GBF1.

Previous reports show significantly decreased FRAP rates for the enzymatically dead GBF1/E794K mutant when expressed alone (Szul et al., 2005). This mutant when expressed in cells causes the dissociation of COPI from the membrane and Golgi disruption (Garcia-Mata et al., 2003). The dominant negative effect of GBF1/E794K expression have been interpreted as GBF1/E794K being stabilized on the membrane by binding to and remaining within an unproductive complex with the endogenous Arf1, thereby sequestering Arf1 from the endogenous GBF1 and preventing Arf1 activation. We show that FRAP recovery for GBF1/E794K when co-expressed with Arf1 is significantly slower relative to wild-type GBF1 co-expressed with Arf1 (t_1/2_ of 19.7 sec versus 7.5 sec for the wild-type GBF1; Table 1). However, when the inverse of the FRAP was plotted on a semi-log, we obtained a single line, suggesting that GBF1/E794K dynamics are limited by a single, relatively slow reaction. This implies that the major contributor to GBF1/E794K overall dynamics are membrane events instead of diffusion. Such events could be due to GBF1/E794K association/reaction with Arf1 or be Arf1-independent.

To distinquish between these possibilities, we examined the dynamics of Arf1 when co-expressed with GBF1/E794K. If GBF1/E794K remains on the membrane in a complex with Arf1, it would be expected that the dynamics of Arf1 would parallel those of GBF1/E794K. However, neither the t_1/2_ nor the type of behavior were the same: while the t_1/2_ of GBF1/E794K recovery was 19.7 sec and its behavior was reaction-limited, the t_1/2_ of Arf1 recovery was 7.2 sec and was diffusion-limited. These results are most consistent with GBF1/E794K dynamics on the membrane being regulated through interactions that may not involve direct Arf1 binding. In this model, GBF1/E794K would induce its disruptive phenotype not by sequestering endogenous Arf1 and preventing its activation as currently accepted, but by saturating membrane-binding sites for the endogenous GBF1. This hypothesis is further validated with the simulation data (Table 2). The ratio of the steady state membrane density, *σ*, to cytosol density, *ρ*, is known to be

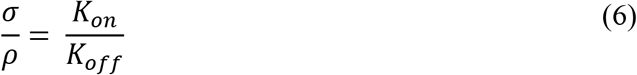

by setting the membrane association rate equal to the membrane dissociation rate. In Table 2, we see *K_on_/K_off_* increases by slightly more than an order of magnitude for the GBF1/E794K coexpressed with Arf1 (*K_on_/K_off_* = 8.67) relative to that of wild-type GBF1 coexpressed with Arf1 (*K_on_/K_off_* = 0.79). This suggests that significantly more GBF1/E794K will be bound to the membrane than wild-type endogenous GBF1, possibly saturating membrane binding sites.

### Implications for Arf1 dynamics on Golgi membranes

Previous FRAP studies reported t_1/2_ of 15 seconds (Presley et al., 2002) and 24 seconds (Szul et al., 2005) for GFP-tagged wild-type Arf1 when expressed alone. Herein, we report a t_1/2_ of 10.3 seconds for Arf1-mCherry. Plotting the inverse of the FRAP data on a semi-log generates two lines, indicating that the dynamics of Arf1 expressed alone are limited by an initial, fast reaction and a latter, slow reaction. We interpret these data to suggest that overproduction of Arf1 without a corresponding increase in the production of GBF1 results in unproductive cycles of Arf1 association and dissociation from the membrane (fast reaction), with only a proportion of Arf1 undergoing productive interaction with the endogenous GBF1, being activated and remaining on the membrane as part of the coating process (slow reaction).

In contrast, when Arf1 is co-expressed with GBF1, the t_1/2_ of its recovery remains similar (10.3 sec versus 12.3 sec), but its dynamics change and are now limited by a single slow process. These data suggest that when overproduction of Arf1 occurs at the same time as a corresponding increase in the production of GBF1, all membrane-associated Arf1 participates in productive interaction with the endogenous and exogenus GBF1, is activated and remains on the membrane in the activated state to facilitate coating (hence only slow reaction is visible). The loss of the fast reaction when ARF1 and GBF1 are co-expressed (perhaps reflecting a more physiological situation) supports a model in which there are more GBF1 on the membrane than Arf1, and thus all Arf1 that associates with the membrane interacts and is activated by GBF1.

This model is supported by the dynamics of Arf1 when co-expressed with the GBF1/7A mutant incapable of binding or activating Arf1. This mutant is expected to be “invisible” to Arf1, but to compete with the endogenous GBF1 for membrane binding sites. Thus, Arf1 was expected to behave similarly to when expressed alone, except that a smaller proportion of Arf1 should be activated and remain on the membrane. Indeed, we observed a slight increase in the FRAP dynamics of Arf1 (t_1/2_ of 7.6 sec versus 10.3 sec for the wild-type GBF1) and two lines when the inverse of the FRAP was plotted on a semi-log. Thus, Arf1 co-expressed with GBF1/7A undergoes fast, ineffective sampling of the membrane and then a reduced level of productive interactions with the endogenous GBF1.

It is important to note that the Arf1 FRAP curves in Figure 3B and 4B, before inverting and graphing on a semi-log plot, look relatively similar. While there are some slight qualitative differences, it would have been impossible to discern how many limiting reactions are present by visual inspection or calculating the t_1/2_. Therefore, when comparing reaction-limited FRAP data, it can be more insightful to use this type of quantitative approach as opposed to mere t_1/2_ value.

The dynamics of GBF1 co-expressed with Arf1/T31N and of Arf1/T31N co-expressed with GBF1 were surprising. Previous studies showed that GBF1 is stabilized in cells when co-expressed with Arf1/T31N (Niu et al., 2005; Szul et al., 2005), and these results were interpreted as GBF1 remaining on the membrane in a complex with the Arf1/T31N mutant. This model implies that GBF1 and Arf1/T31N should show similar FRAP behaviors on the membrane and be reaction-limited. The FRAP dynamics of GBF1 co-expressed with Arf1/T31N (t_1/2_ of 6.7 sec) were different from those of Arf1/T31N (t_1/2_ of 2.9 sec), and the dynamics were diffusion-limited. This suggests that even if GBF1 and Arf1/T31N interact on the membrane, such interactions are so fast as to be “invisible” in the overall dynamics. Alternatively, GBF1 and Arf1/T31N might participate in limited interaction/reaction events on the membrane. The latter interpretation is supported by the limited co-localization of GBF1 and Arf1/T31N in cells (Chun et al., 2008; Szul et al., 2005). These data suggest that Arf1/T31N would induce its disruptive phenotype not by sequestering endogenous GBF1 and preventing the activation of endogenous Arf1 as currently accepted, but by reducing the membrane concentration of the endogenous Arf1.

#### Role of Simulations in Analyzing GBF1 and Arf1 dynamics

the simulations quantify how much endogenous proteins can contribute to kinetic data, especially when overexpressing certain types of mutants, and provide insight into possible mechanisms through which protein behavior is influenced by the co-expressed proteins. Our simulations generally fit well with our experimental data, with the exception of Arf1 dynamics when co-expressed with the GBF1/E794K mutant. In this case, both the endogenous GBF1 and the GBF1/E794K are present in the cell and Arf1 dynamics can be impacted by both GBF1 species, resulting in dynamics that reflect the composite of those distinct behaviors. Our inability to make the simulation fit the experimental FRAP data indicates a large contribution of the endogenous GBF1 to Arf1 dynamics. Future studies can avoid the contribution of the endogenous proteins when analyzing kinetic data by knocking down the endogenous protein and assuring that only the exogenously produced protein influences the dynamics.

Coating is a compilation of distinct subprocesses linked to each other temporally and causally, and to understand the overall process, we need to know how one component influences the behavior of other components within the coating pathway. Understanding the overall order of the distinct subprocesses and the relative impact of each subprocess on subsequent events will require additional studies to incorporate the dynamics of additional molecules within the coating pathway including ArfGAP1 and coatomer. Ultimately, incorporation of additional Arf1 and GBF1 interactors will provide a network-level understanding of the coating process.

## MATERIALS AND METHODS

### Plasmids

mCherry-tagged wild-type Arf1 and Arf1/T31N constructs were a generous gifts from Dr. Paul Melançon (University of Alberta, Alberta, Canada). Green fluorescent protein (GFP)-tagged wild-type GBF1 and GBF1/E794K have been previously described (Garcia-Mata et al., 2003). GFP-tagged GBF1/7A has been previously described (Lowery et al., 2011).

### Cell Culture and Transfection

HeLa cells were grown in DMEM media (Corning) supplemented with 10% Fetal Bovine Serum and 1% penicillin/streptomycin (Corning) at 37°C in 5% CO_2_. Cells were grown on a 25 mm diameter glass cover slips of thickness #1.5 (Electron Microscopy Sciences) for ~24 hours pre-transfection till ~70% confluent. 1.0 μg of single plasmid DNA or 0.5 μg of each plasmid DNA for co-transfection experiments was added to 100 μL of RPMI-1640 with L-Glutamine (Corning) and 5 μL of TransIT-LT1 (Mirus), and the transfection was performed as per manufacturer’s directions. Cells were grown for 24 hours prior to FRAP.

### Fluorescence Recovery after Photobleaching (FRAP)

the Nikon A1R-HD25 microscope at the University of Alabama at Birmingham High Resolution Imaging Facility Service Center was used for FRAP analysis. The cells were placed inside a silicon sealed chamber at 37°C and 5% CO_2_. For GFP-tagged constructs, a 488-nm laser beam was used to photobleach an area of 2 μm on the Golgi. For mCherry-tagged constructs, the laser wavelength was set to 561 nm. Frames were captured every 1s, and the data were averaged over 9-12 cells. For GBF1/E794K co-expressed with Arf1, 12 cells were averaged. For GBF1 co expressed with Arf, 11 cells were averaged. For GBF1/7A co-expressed with Arf1, 9 cells were averaged. For GBF1 co-expressed with Arf1/T31N, 10 cells were averaged.

All FRAP data are normalized and corrected for non-specific photobleaching by subtracting the background intensity from the experimental and reference regions of interest (ROI) and taking the ratio, respectively. The experimental ROI is the area of the Golgi being bleached, while the reference ROI is in the cytosol, far from the experimental region, to correct for any photobleaching. The photobleached corrected data is collected by dividing the experimental ROI by the reference ROI for each frame. Then, data are divided by the intensity immediately before bleaching to obtain a normalized recovery plot.

### Monte Carlo Simulation

the simulation is written in Fortran 95 and uses a discrete space, continuous time random walk for all molecular species (Kosztolowicz, 2015). The diffusion is simulated as jump processes with a fixed distance approximately the size of the molecules. The jumps are considered as Poison processes with a given mean transition time. The event driven algorithm based on binary tree calendar is adopted for event scheduling to increase computational efficiency. In total, there are 14 parameters capturing the diffusion, membrane association/dissociation, and reaction rates for all molecular species.

### Plotting FRAP Data on Semi-log

As with any data analysis workflow, there is always a push-pull relationship between developing a method that is convenient to use and maintaining scientific rigor. The semi-log method for FRAP data is not optimal to assess the behavior of proteins limited by highly nonlinear reactions of second order or higher. Nonlinear differential equations are often impossible to analytically solve, and simple equation (such as Eq. 2 or Eq. 4) will not fit these data. If this occurs, it would be recommended to use other metrics (such as the t_1/2_ and the immobile phase) to analyze the FRAP data. Considering cellular processes are usually non-linear, it may seem the semi-log method will almost always fail and therefore be useless. However, since the system is in steady state prior to the photobleaching, the nonlinear reaction-diffusion equations for molecules in the FRAP experiment can sometimes be simplified to linear equations.

Generally, it is assumed that the concentrations of all molecular species in a reaction-diffusion PDE are time dependent. This assumption is invalid for FRAP, because the system was in steady state before bleaching, and remains so during the subsequent recovery. To accurately solve the reaction-diffusion equation for the fluorescent Arf1-GDP molecules, the total number of GBF1 molecules in the cell should be treated as a constant (assuming photobleaching does not dramatically affect the reactivity of GBF1). Hence, any reaction term in the Arf1-GDP reaction-diffusion equation that involves the product of the concentration of Arf1-GDP and GBF1 would become linear, since the concentration of GBF1 is constant in that specific equation (note, the opposite will be true for the reaction-diffusion equation of GBF1). Hence, the semi-log plot will still appear linear for molecules involved in nonlinear reactions when the nonlinearity arises from the product of the concentration of substrates. This simplification also eliminates most, if not all, the coupling terms between differential equations, which may make the reaction-diffusion ODEs analytically solvable. Hence, the semi-log method should be helpful for a large set of limiting reactions. In this report, we show that the semi-log method can be used to describe the limiting processes of Arf1 and GBF1 diffusion and association with Golgi membranes.

## ACKNOWLEDGMENTS

We wish to thank Drs. Mayorga and Kahn for their important insights and suggestions during the course of this study. This work was supported by grants from the NSF (MCB-1615607 to ES) and the NIH (R01GM122802 to ES). The High Resolution Imaging Facility Service Center is an institutional core at UAB supported by the Office of the Vice President of Research and Development and the following grants: Cancer Center Support Grant P30 CA013148 and Rheumatic Disease Core Center P30 AR048311.

## Abbreviations

GBF1: Golgi Brefeldin A Resistant Guanine Nucleotide Exchange Factor 1
ARF1: ADP-ribosylation factor 1
FRAP: fluorescence-recovery-after-photobleaching

