## Supplemental Tables for "Modeling the dynamic behaviors of the COPI vesicle formation regulators, the small GTPase Arf1 and its activating Sec7 guanine nucleotide exchange factor GBF1 on Golgi membranes"

Supplemental Table 1: Examined combinations of GBF1 and Arf1 constructs.

| GBF1 Protein | Arf1 Protein |
|---|---|
| GBF1 WT GFP | - |
| - | Arf1 WT mCherry |
| GBF1 WT GFP | Arf1 WT mCherry |
| GBF1 E794K GFP | Arf1 WT mCherry |
| GBF1 7A GFP | Arf1 WT mCherry |
| GBF1 WT GFP | Arf1 T31N mCherry |

Supplemental Table 1: The table lists all the combinations of components analyzed in this study. A “-“ means no exogenous protein of that species was introduced into cells.

Supplemental Table 2: P values for $t_{1/2}$ of GBF1 dynamics from FRAP data.

| Experimental Condition | Mean ± STD | P-values | | | | | | | | | |
|---|---|---|---|---|---|---|---|---|---|---|---|
| GBF1 WT (without Arf1 coexpression) | 8.0 ± 2.5 | 8.3E-4 | 9.0E-1 | 4.1E-4 | 4.4E-3 | | | | | | |
| GBF1 WT (with Arf1 WT coexpression) | 4.2 ± 1.5 | ** | | | | 2.7E-3 | 4.4E-5 | 2.8E-1 | | | |
| GBF1 WT (with Arf1/T31N coexpression) | 7.8 ± 2.8 | | | | | ** | | | 3.7E-4 | 1.2E-2 | |
| GBF1/E794K (with Arf1 WT coexpression) | 20.6 ± 8.9 | | | *** | | | **** | | *** | | 6.7E-5 |
| GBF1/7A (with Arf1 WT coexpression) | 4.9 ± 1.4 | | | | ** | | | | | * | *** |

Supplemental Table 2: The table lists the mean ± standard deviation (STD) $t_{1/2}$ values for GFP-GBF1 constructs expressed alone or co-expressed with Arf1-mCherry constructs. P-values are calculated with a two-tailed t-test and using $\alpha = 0.05$. To reduce space, scientific notation (i.e. $3 \times 10^{-5}$) is abbreviated with an E (i.e. 3E-5). With five experimental conditions, there are a total of ten p-values calculated. Each pair of comparisons is denoted by similar color. Light green corresponds to the results of the t-test analysis between GBF1 expressed alone and GBF1 co-expressed with Arf1. Light blue corresponds to the results of the t-test analysis between GBF1 expressed alone and GBF1 co-expressed with Arf1/T31N. Light tan corresponds to the results of the t-test analysis between GBF1 expressed alone and GBF1/E794K co-expressed with Arf1. Light gray corresponds to the results of the t-test analysis between GBF1 expressed alone and GBF1/7A co-expressed with Arf1. Light pink corresponds to the results of the t-test analysis between GBF1 co-expressed with Arf1 and GBF1 co-expressed with Arf1/T31N. Blue corresponds to the results of the t-test analysis between GBF1 co-expressed with Arf1 and GBF1/E794K co-expressed with Arf1. Green corresponds to the results of the t-test analysis between GBF1 co-expressed with Arf1 and GBF1/7A co-expressed with Arf1. Yellow corresponds to the results of the t-test analysis between GBF1 co-expressed with Arf1/T31N and GBF1/E794K co-expressed with Arf1. Dark gray corresponds to the results of the t-test analysis between GBF1 co-expressed with Arf1/T31N and GBF1/7A co-expressed with Arf1. Dark pink corresponds to the results of the t-test analysis between GBF1/E794K co-expressed with Arf1 and GBF1/7A co-expressed with Arf1. The top box shows the p-value and the bottom box demonstrates the level of significance. No asterisks are used for p-values greater than 5E-2, one asterisk (*) is used for p-values less than 5E-2, two asterisks are used for p-values less than 5E-3, three asterisks are used for p-values less than 5E-4, and four asterisks indicate p-values less than 5E-5.

Supplemental Table 3: P values for $t_{1/2}$ of Arf1 dynamics from FRAP data.

| Experimental Condition | Mean ± STD | P-values | | | | | | | | | |
|---|---|---|---|---|---|---|---|---|---|---|---|
| Arf1 WT (without GBF1 WT coexpression) | 9.3 ± 2.7 | 2.6E-1 | 5.5E-4 | 9.2E-1 | 9.9E-1 | | | | | | |
| Arf1 WT (with GBF1 WT coexpression) | 11.0 ± 3.5 | | | | | 1.0E-5 | 2.0E-1 | 4.6E-1 | | | |
| Arf1/T31N (with GBF1 WT coexpression) | 3.2 ± 1.5 | | ** | | | **** | | | 3.9E-5 | 1.9E-2 | |
| Arf1 WT (with GBF1/E794K coexpression) | 9.1 ± 3.3 | | | | | | | | **** | | 9.6E-1 |
| Arf1 WT (with GBF1/7A coexpression) | 9.2 ± 6.1 | | | | | | | | | * | |

Supplemental Table 3: The table lists the mean ± standard deviation (STD) $t_{1/2}$ values for Arf1-mCherry constructs co-expressed with GFP-GBF1 constructs. P-values are calculated with a two-tailed t-test and using $\alpha = 0.05$. To reduce space, scientific notation (i.e. $3 \times 10^{-5}$) is abbreviated with an E (i.e. 3E-5). With five experimental conditions, there are a total of ten p-values calculated. Each pair of comparisons is denoted by similar color. Light green corresponds to the results of the t-test analysis between Arf1 expressed alone and Arf1 co-expressed with GBF1. Light blue corresponds to the results of the t-test analysis between Arf1 expressed alone and Arf1/T31N co-expressed with GBF1.Tan corresponds to the results of the t-test analysis between Arf1 expressed alone and Arf1 co-expressed with GBF1/E794K. Light gray corresponds to the results of the t-test analysis between Arf1 expressed alone and Arf1 co-expressed with GBF1/7A. Light pink corresponds to the results of the t-test analysis between Arf1 co-expressed with GBF1 and Arf1/T31N co-expressed with GBF1. Blue corresponds to the results of the t-test analysis between Arf1 co-expressed with GBF1 and Arf1 co-expressed with GBF1/E794K. Green corresponds to the results of the t-test analysis between Arf1 co-expressed with GBF1 and Arf1 co-expressed with GBF1/7A. Yellow corresponds to the results of the t-test analysis between Arf1/T31N co-expressed with GBF1 and Arf1 co-expressed with GBF1/E794K. Dark gray corresponds to the results of the t-test analysis between Arf1/T31N co-expressed with GBF1 and Arf1 co-expressed with GBF1/7A. Dark pink corresponds to the results of the t-test analysis between Arf1 co-expressed with GBF1/E794K and Arf1 co-expressed with GBF1/7A. The top box shows the p-value and the bottom box demonstrates the level of significance. No asterisks are used for p-values greater than 5E-2, one asterisk (*) is used for p-values less than 5E-2, two asterisks are used for p-values less than 5E-3, three asterisks are used for p-values less than 5E-4, and four asterisks indicate p-values less than 5E-5.
